# Dispersal ability of *Neophilaenus campestris*, a vector of *Xylella fastidiosa*, from olive groves to over-summering hosts

**DOI:** 10.1101/2020.03.17.995266

**Authors:** C. Lago, M. Morente, D. De las Heras-Bravo, A. Marti Campoy, F. Rodriguez-Ballester, M. Plaza, A. Moreno, A. Fereres

## Abstract

*Neophilaenus campestris* is one of the recently identified spittlebugs (Hemiptera: Cercopoidea) able to transmit *Xylella fastidiosa* to olive trees. Considering its vector ability and the wide distribution of this species in Spain, *N. campestris* should be considered a serious threat to key crops that are vital for Spanish agriculture such as olive, almonds and grapevines. Migration and dispersal abilities of insect vectors have profound implications in the spread of vector-borne diseases. Thus, knowledge on the dispersal ability of *N. campestris* is essential to model, predict and limit the spread of the diseases caused by *X. fastidiosa*. A mark-release-recapture technique was developed to track between-field movements of *N. campestris* during its late spring migration from the ground cover grasses within olive groves to sheltered areas dominated by pine trees. An indoor assay showed that the fluorescent dust used for marking did not affect the survival nor the flying ability of *N. campestris*. Spittlebug adults captured in olive groves at Los Santos de la Humosa (Madrid, Spain) during late spring, 2019 were dusted with four fluorescent colours and released in four different locations. Six recapture samplings were performed 23 to 42 days after release in 12 different sites located within a maximum distance of 2.8 km from the release point. Results indicated that *N. campestris* was able to disperse a maximum distance of 2473 m in 35 days from the olive groves to areas dominated by pine trees. Furthermore, our flight mill studies also showed that *N. campestris* was able to fly long distances, reaching almost 1.4 km in an 82 minutes’ single flight.

Moreover, we carried out a survey of directional movement of potential vectors of *X. fastidiosa* in an olive grove located in Villa del Prado (Madrid). We used yellow sticky bands, a Malaise trap and a vertical yellow sticky net to assess the directional movement from olive groves to surrounding managed and unmanaged areas. The captures obtained in the yellow sticky bands showed that spittlebugs dispersal from the olive grove to surrounding vegetation matched with the time when the ground cover dried out. The highest number of spittlebugs was captured in the border between the olive grove and a vineyard close by.

Altogether, our findings suggest that eradication measures by rooting-up *X. fastidiosa*-infected and non-infected trees in a radius of 100 m are of limited value because vectors are able to disperse rapidly over distances much longer than expected.

## 1. INTRODUCTION

*Xyllela fastidiosa* Wells (1987) is a vector-borne plant pathogenic bacterium native to the Americas, which has been recently detected in Europe (EFSA, 2019). The bacterium is responsible for severe diseases of several economically important crops such as olive, almond, grapevines and citrus (Hopkins, 1989; Saponari et al., 2013). In Europe, the bacterium was firstly detected in 2013 in Apulia, southern Italy, (Saponari et al., 2013) where it is responsible for the Olive Quick Decline Syndrome (OQDS), a disease that killed more than a million olive trees in this region (Martelli et al., 2016; Saponari et al., 2017; EPPO, 2018). After the detection of *X. fastidiosa* in Italy, the European Union developed a large-scale surveying plan focused on the detection of the bacterium in different economically important crops throughout Europe. As a result, the pathogen was detected in France, Germany, Portugal and Spain (EFSA, 2019). Very recently, *X. fastidiosa* was also detected in Northern Israel (EPPO, 2019). In Spain, *X. fastidiosa* was first detected in the Balearic Islands in 2016 on cherry, and months later on almond, wild and cultivated olive and grapevines among other crops. In 2017, the bacterium was also found in the Valle de Guadalest (Alicante), on almond trees (MAPA. GOB, 2019) and the disease has now spread to a wide area close to 140,000 ha (Generalitat Valenciana, 2019). *X. fastidiosa* was also found on olives in Villarejo de Salvanés (Madrid, Spain) and *Polygala myrtifolia* in Almeria, Spain, although these infections were officially declared eradicated.

*X. fastidiosa* is transmitted to plants exclusively by xylem-sap feeding insects (Frazier 1965). Insects that are exclusively xylem-sap feeders, thus putative vectors of the fastidious bacterium, belong to the Order Hemiptera, suborder Cicadomorpha, Superfamilies Cercopoidea (spittlebugs or froghoppers) and Cicadoidea (cicadas), as well as to the Family Cicadellidae, Subfamily Cicadellinae (sharpshooters) (Novotny & Wilson, 1997; Redak et al., 2004; Almeida et al., 2005; Krugner et al., 2019). While sharpshooters are overall scarce in Europe, spittlebugs are much more abundant, thus they are considered as the main potential vectors of *X. fastidiosa* in the European continent (Cornara et al., 2019; Jacques et al., 2019). The meadow spittlebug, *Philaenus spumarius* L. (1758) (Hemiptera: Aphrophoridae) was identified as the main vector of *X. fastidiosa* in the olive groves of southern Italy (Cornara et al., 2016; Cornara et al., 2017a). Moreover, the spittlebugs *Neophilaenus campestris* Fallen (1805) and *Philaenus italosignus* Drosopoulos & Remane (2000) have been found to transmit *X. fastidiosa* to olive and other plants under experimental conditions, although less efficiently than *P. spumarius* (Cavalieri et al. 2019). Recently, Cornara el al. (2020b) found that the main European cicada species, *Cicada orni,* is unable to transmit the ST53 strain of *X. fastidiosa* to olive plants.

Migratory journeys and dispersal abilities have profound implications in the spread of vector-borne diseases (Irwin & Thresh 1988; Chapman et al., 2015; Fereres et al., 2017). Adults of *P. spumarius* and related species are able to actively disperse, with a migratory behaviour been observed by several authors (Weaver, 1951; Weaver & King, 1954; Lavigne, 1959; Halkka et al., 1967; Halkka et al., 1971; Cornara et al., 2018; Bodino et al. 2019). *P. spumarius* and *N. campestris* adults spend most of their life cycle on the ground vegetation (Mazzoni, 2005; Dongiovanni, 2018; Morente et al., 2018a). The first migration seems to happen in summer (Weaver & King, 1954; Waloff, 1973). It has been observed that they displace from the ground when the grasses dry out, to woody hosts and evergreen or deciduous plant species (Lopes et al., 2014; Cornara et al., 2016; Morente et al., 2018a; Antonatos et al., 2019). Later in the fall, spittlebugs leave their woody hosts to lay their eggs after the first rains on ground cover vegetation and plant debris present in olive groves (Cruaud et al., 2018; Morente et al., 2018a; Antonatos et al., 2019). Morente et al., (2018a) and Lopes et al., (2014) have reported that *N. campestris* are abundant in pine trees (*Pinus halepensis*) in the summer months in continental Spain. This suggests that pine trees could be an over-summering host plant exploited by *N. campestris* as a shelter when the grasses dry out. Despite spittlebugs spend most of the time on the ground vegetation, it has been proposed that they may play an important role in *X. fastidiosa* transmission when they displace from grasses in the late spring to feed on woody hosts (Almeida, 2016; Morente et al., 2018a). In the process of selecting their over-summering host they can settle and feed on woody crops such as almond and grapevine where they can transmit the disease (Purcell, 1980). Indeed, since the process of transmission of *X. fastidiosa* may occur in few minutes (Cornara et al., 2020a), non-colonizing spittlebug species may have an impact on disease epidemiology. This could be the case of *N. campestris* that is frequently found in ground cover vegetation in olive groves but is rarely found feeding on the olive tree canopy (Morente et al., 2018a; Bodino et al., 2019).

Weaver and King (1954) observed that marked *P. spumarius* travelled more than 30 m in a single flight, and moved as much as 100 m within 24 hours from the release point. The same authors also observed that *P. spumarius* adults mainly fly at a height of 15 to 70 cm above the ground. In contrast, Freeman (1945) collected *P. spumarius* and *N. lineatus* at 84 m above ground and Reynolds et al. (2017) reported captures of *N. lineatus* at 200 m above ground suggesting that they can reach much higher altitudes. Migrating insects are strongly influenced by the planetary boundary layer, the lowest part of the troposphere, which is defined by turbulent convective air motions and stable laminar air currents (Caughey, 1984; Drake & Farrow, 1988; Isard et al., 1990). Near the ground, the speed of flying insects is higher than the wind speed, so insects are capable to intentionally displace to, a specific zone in the atmosphere defined as the flight boundary layer (Southwood, 1962; Taylor, 1974; Isard et al., 1990). When insects travel above the flight boundary layer, they can reach a stable part of the planetary boundary layer, where the wind speed is maximum usually at few hundred meters in height. Thus, insects reaching the planetary boundary layer can be transported by these winds commonly known as low-level jet winds (Gerhardt, 1962; Drake & Farrow, 1988). Several studies show that even the weakly flying insects can be transported long distances due to low-level jet winds (Pienkowski & Medler, 1964; Drake, 1985; Wallin & Loonan, 1971; Sedlacek & Freytag, 1986; Zhu et al., 2006). Thus, all the captures in altitude of spittlebugs previously reported by several authors strongly suggest that *X. fastidiosa* vectors may perform long distance migrations.

Studying dispersal patterns and insect migration behaviour requires insect tracking in the field, which can be challenging due to their small size and general lack of specific return-migration sites (Chapman et al., 2015). Nevertheless, a combination of several methods can improve our knowledge on the movement and dispersal behaviour of the vectors of *X. fastidiosa* (Purcell et al., 1994). Mark-release-recapture studies using multiple types of markers have been used since 1920s to understand the movement of insects in the field (Hagler & Jackson, 2001; Hagler, 2019). Fluorescent dusts are one of the most extended markers used for mark-release-recapture tests (Byrne et al., 1996; Prasifka et al., 1999; Hagler & Jackson, 2001; Miranda et al., 2018). This technique has been largely used to study the movement of important agricultural insect pests, including the leafhopper *Scaphoideus titanus Ball*, which is the vector of the Flavescence dorée plant disease, or American vectors of *X. fastidiosa* such as *Homalodisca vitripennis* (Germar 1821) (Coviella et al., 2006; Northfield et al., in 2009; Lessio et al., 2014). Additionally, other techniques based on the interception of the insect’ displacement provide information about the movement of flying insects across habitat boundaries (Stewart, 2002).

Alternatively, laboratory-based flight mills have been used since the 1950s to generate knowledge on the flight behaviour of several order of insects: Orthoptera (Krogh & Weis-Fogh, 1952), Lepidoptera (Guo, 2020), Coleoptera (Ávalos, 2014; Yu et al., 2019), Hemiptera (Taylor, 1992; Riley et al., 1997, Martini et al., 2015) or Diptera (Somerville, 2019). Despite the broad variety of flight mill designs, they are all based on the same principle: an insect is attached to an arm, which is connected to a stand, then the insect flies describing a circular trajectory, allowing continuous measurement of flight parameters (Minter, 2018). This tool has been applied to study the dispersal ability of serious insect pests, such as the red palm weevil (Avalos et al., 2014) or the western corn rootworm (Yu et al., 2019). An interesting application of flight mills is to describe how a plant pathogen may modify the flying ability of its vector such as the Asian citrus psyllid when infected with *Candidatus liberibacter* asiaticus (Martini et al., 2015). Moreover, flight mills are commonly used to describe how abiotic factors (humidity, temperature) or biotic factors (age, sex, matted status) influence insect displacement (Riley et al., 1997; Zhang et al., 2008; Cheng et al., 2012; Jones et al., 2015). The vast literature of studies using flight mills probe that this tool, contribute to understand the flight behaviour of important insect pests.

One of the *X. fastidiosa* eradication measures that are mandatory by the European Commission (EU 2015/789, Article 6) consists in up-rooting the infected plants and the entire host plants in a radius of 100 m, regardless their health status. This strategy is mainly based on the Weaver & King study (1954), which describes that *P. spumarius* can travel as much as 100 m in 24 hours. However, to date there is no precise information on how far spittlebugs can fly when they migrate from cultivated fields to their over-summering hosts.

Therefore, the main aim of this work was to understand spittlebug dispersal dynamics by combining different techniques: (1) flight mill assays capture-mark-release-recapture assay and (3) directional movement study using interception traps.

## 2. MATERIAL AND METHODS

An indoor study to assess the persistence of fluorescent dusts (Day-Glo Color Corp. Cleveland, OH, USA) and its effect on the survivorship and flight ability of *N. campestris* was conducted before the mark-release-recapture assay. This type of fluorescent dusts has been used for many years to mark insects and study their dispersal ability in the field (Stern et al., 1965). *N. campestris* adults were collected by sweep net in Los Santos de la Humosa (Madrid, Spain) in late spring 2019; the location was the same where the mark-release-recapture assay was performed. Spittlebugs collected were identified according to Ribaut (1952), Ossiannillsson (1981), Della Giustina (1989), Holzinger et al. (2003) and Mozaffarian & Wilson (2016). Insects collected were caged on *Bromus madritensis* during 3 days for acclimation in the greenhouse facilities at ICA-CSIC, Madrid, Spain.

### 2.1. Persistence of fluorescent dusts and their effect on the survival of *Neophilaenus campestris*

To assess the effect of the fluorescent dusts on the survival of *N. campestris*, 200 individuals were randomly split in 5 groups: a dusted group which included one of each of the following colours: pink, blue, yellow and orange and an undusted control group. We used 2.8 mg of fluorescent dust per 40 insects for each of the dusted groups. The dust was first introduced in the falcon tube and then 40 adult insects/tube were released. Then, the falcon tube was gently shaken to allow the dust to cover most of the insect’s body. Then, each group of insects was released in a cage (40 adults per cage) containing 4-week old potted *B. madritensis* plants (plants grown in a climatic chamber at 24:18°C of temperature and photoperiod 14:10). The number of alive and dead insects on each cage and the persistence of the dust on the insect’s body were recorded twice a week during 35 days. A 4-level scale of dust coverage was established, in relation to the intensity of the fluorescence on the insects. 1) completely dusted, 2) less dust but visible at naked eye, 3) fluorescence not visible at naked eye but visible by using UV light, 4) undusted. The assay was conducted in a greenhouse at ICA-CSIC (Madrid) (Temperature (mean ± SE): 22.28 ± 0.23ºC; Max 40.06ºC; Min 7.99ºC. RH (mean ± SE): 54.64 ± 0.61%; Max: 99.31%; Min 19.95%). The plants were replaced every week to keep optimal conditions for insect rearing. We performed a two-sample Cox proportional hazards model to determine whether the colour of the fluorescent dust affected the survival of adults (R Core Team, 2019).

### 2.2. Effect of fluorescent dusts on the flight behaviour of *Neophilaenus campestris*

A commercial flight mill device (Insect FlyteMill, Crist Instruments, Hagerstown, MD, USA) with some adaptations to reduce friction and facilitate the flight of small insects was used to evaluate the effect of the dust on the flight potential of *N. campestris*. Flight mill recordings were taken 1-3 days after the insects were dusted with fluorescent dust using the same methodology for marking and the same 5 experimental groups (4 dusted and one undusted) described above. Individuals were exposed to greenhouse conditions (T (Mean ± SE): 22.28 ± 0.23ºC; Max 40.06ºC; Min 7.99ºC. RH (Mean ± SE): 54.64 ± 0.61%; Max: 99.31%; Min 19.95%) until the experiments were started. Experiments were carried out in the laboratory under controlled conditions: temperature (24±1Cº), artificial fluorescent light (10 μE m−2 s−1) and humidity (25-55%). The experiments took place from 9 AM to 18 PM. Insects were first anesthetized by applying CO_2_ during 5 seconds. Then, they were glued to a pinhead by the pronotum using a small drop of adhesive (Hot melt glue, NV98591 Nivel, Leganes, Madrid, Spain). Then, the insects were placed on one side of the flight mill’s arm (29.6 cm) with a suitable counter balance on the opposite side of the arm to make them fly in a circular trajectory. Insects that did not start to fly after 15 minutes were removed and discarded. The flight activity was recorded until the insect stopped flying for a time interval longer than 15 minutes.

The data collected by the flight mill device were the following: the distance flown (m), the total flight duration (s) and the flight speed (m/s). A specific “mill_recorder” computer-based software and hardware device recorded the data and the “mill_processor” software calculated the flight descriptors (both developed by Marti-Campoy & Rodriguez-Ballester at the ITACA-Universitat Politècnica de València, Valencia, Spain). The flight potential was evaluated according to the following flight descriptors: (1) Flight incidence: the ability of a given insect to perform a flight (Yes/No); (2) Number of flights: a new flight in the recording was assumed when an insect spent more than 20 seconds to complete one turn and until a 15 minutes’ lag; Total distance travelled: sum of the distance covered by all flights; (4) Total duration: sum of the duration of all flights; (5) Average speed: mean of the speed of each individual flight. The maximum distance travelled, flight duration and average speed were also recorded. A total of 89 individuals were tested until completion of 10 flight recordings for each of the 5 experimental treatments-dusted and undusted-(50 recordings in total).

We compared the effect of the dust in the flight behaviour of the 5 different experimental groups for each of the flight descriptors (dependent variables) by an ANOVA or a Kruskal-Wallis test depending on the distribution of the data (Gaussian or non-Gaussian distribution, respectively). The probability of performing or not a flight was tested using a Chi-squared test. Data of non-flyers was not considered for the analysis. Statistical analysis was performed using the SPSS software (IBM SPSS Statistics 25).

### 2.3. Mass-Mark-Recapture assay (MMR)

The study was conducted at Los Santos de la Humosa, Eastern Madrid (Spain) (40° 30’ 04.08’’ N 3° 15.25’ 58’’ W, 850 m). We used 4 different colours (pink, blue, yellow and orange) for marking insects that were released in 4 different olive groves separated 200 m from each other (one colour per grove). The different colours were used to identify the distance travelled from each of the 4 release points to the recapture sites. The insect releases were carried out in olive groves with abundant ground cover vegetation, mainly dominated by grasses (Poaceae) (Figure 1, Annex 1). The selection of the recapture sites was based on the presence of perennial natural woody vegetation, which included known host species of *N. campestris* and other spittlebug species present in the area such as the vector of *X. fastidiosa P. spumarius* and the potential vector *Lepyronia coleoptrata* (Lopes et al., 2014; Morente et al., 2018a). Thus, recapture sampling procedure was performed in 12 different sites where the dominant vegetation was *Pinus halepensis*, *P. pinea*, *Quercus coccifera*, *Q. faginea*, *Retama sphaerocarpa*, *Foeniculum vulgare, Eryngium campestre* and *Prunus dulcis* (Figure 1). Recapture points were located at different distances, being the minimum distance 94 m and the maximum of 2754 m from the most distant release point (Figure 1). The first spittlebug capture-mark-release procedure was carried out on 23rd May 2019 following a methodology similar to the one described by Nakata (2008). Adult individuals were captured by a sweep net from the ground cover vegetation in the four olive groves mentioned above and stored in 50 ml conical falcon tubes. Individuals captured were dusted in groups of 100 insects per falcon tube. Thus, the insects were introduced in a falcon tube that contained 7 mg of fluorescent dust (Day-Glo Color Corp., Cleveland, OH, USA). The same procedure was repeated with each of the 4 different colours. The dusted spittlebugs were released on the green ground cover of each olive grove. The first recapture event was carried out on 12th June 2019, 20 days after the release date and matching with the senescence of the ground cover vegetation. In total we performed the following five recaptures: 12th, 18th, 19th, 20th, 27th June and 5th July. Because the fluorescent dust was not visible at naked eye, insects were recaptured by sweep net and caged on *B. madritensis* plants and transferred to the laboratory. We assessed the presence of fluorescent dust on the body of every individual by using a UV lamp 13W (Halotec F6T5/BLB, Koala Components, Torrent (Valencia), Spain).

**FIGURE 1.**
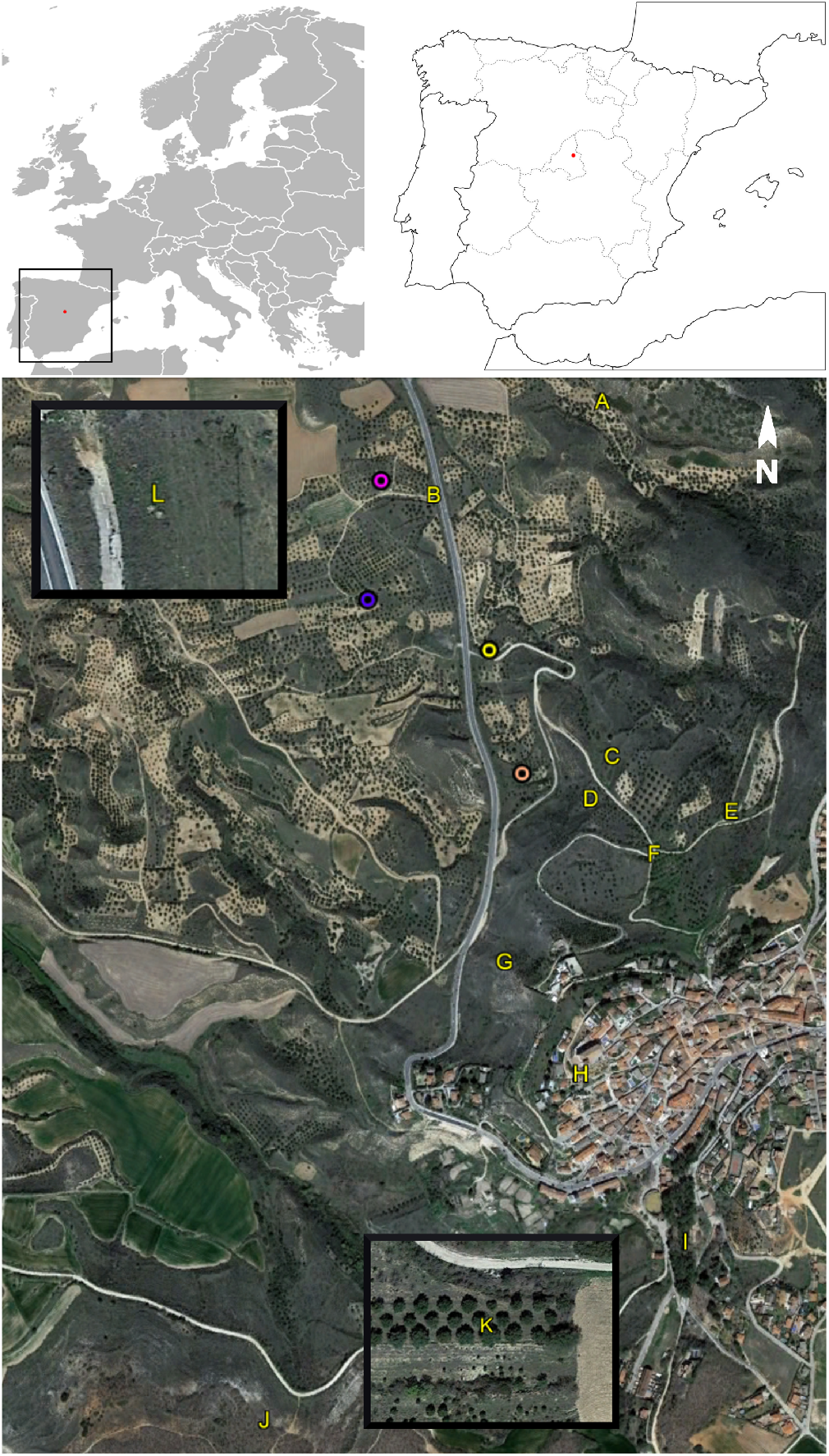
Mark-release-recapture study zone. 1) Coloured circles: release points (pink, blue, yellow and orange) 2) Letters: recapture points. A: *Quercus coccifera*; B: *Foeniculum vulgare*, *Eryngium campestre* and Asteraceae; C: *Retama sphaerocarpa* and *E. campestre*; D: *Pinus halepensis*; E: *Prunus dulcis*; F: *P. dulcis*; G, H and I: *Pinus halepensis*; J: *Q. faginea* K: *P. halepensis* and *P. pinea*; L: *Foeniculum vulgare* and *Retama sphaerocarpa*. 3) Points L and K shown in the upper left and lower right corner, respectively are out of the map scale because they were located too far away from the release points. Google. (n.d.). Los Santos de la Humosa. Retrieved from https://www.google.es/maps/place/28817+Los+Santos+de+la+Humosa,+Madrid/@40.5020998,-

Regarding the high adherence of the fluorescence dust we carried out several precautionary measures in order to avoid the contamination of the individuals recaptured. First, we replaced every day all materials used in the marking process (i.e. plastic bags and Falcon tubes). Moreover, we replaced the Falcon tubes every day of recapture and the insect mouth aspirators were inspected every day checked under UV light looking for fluorescence traces. Furthermore, the individuals recaptured at the field were stored in groups of 50 individuals in falcon tubes and, later, caged on *B. madritensis* plants (one cage per location of recapture and date), for transportation to the laboratory. Then, all the recaptured individuals were checked under UV light and screened for the presence of fluorescent dust in the insect’s body. We considered a marked spittlebug those that showed clear trace of fluorescent dust (Figure 2).

**FIGURE 2.**
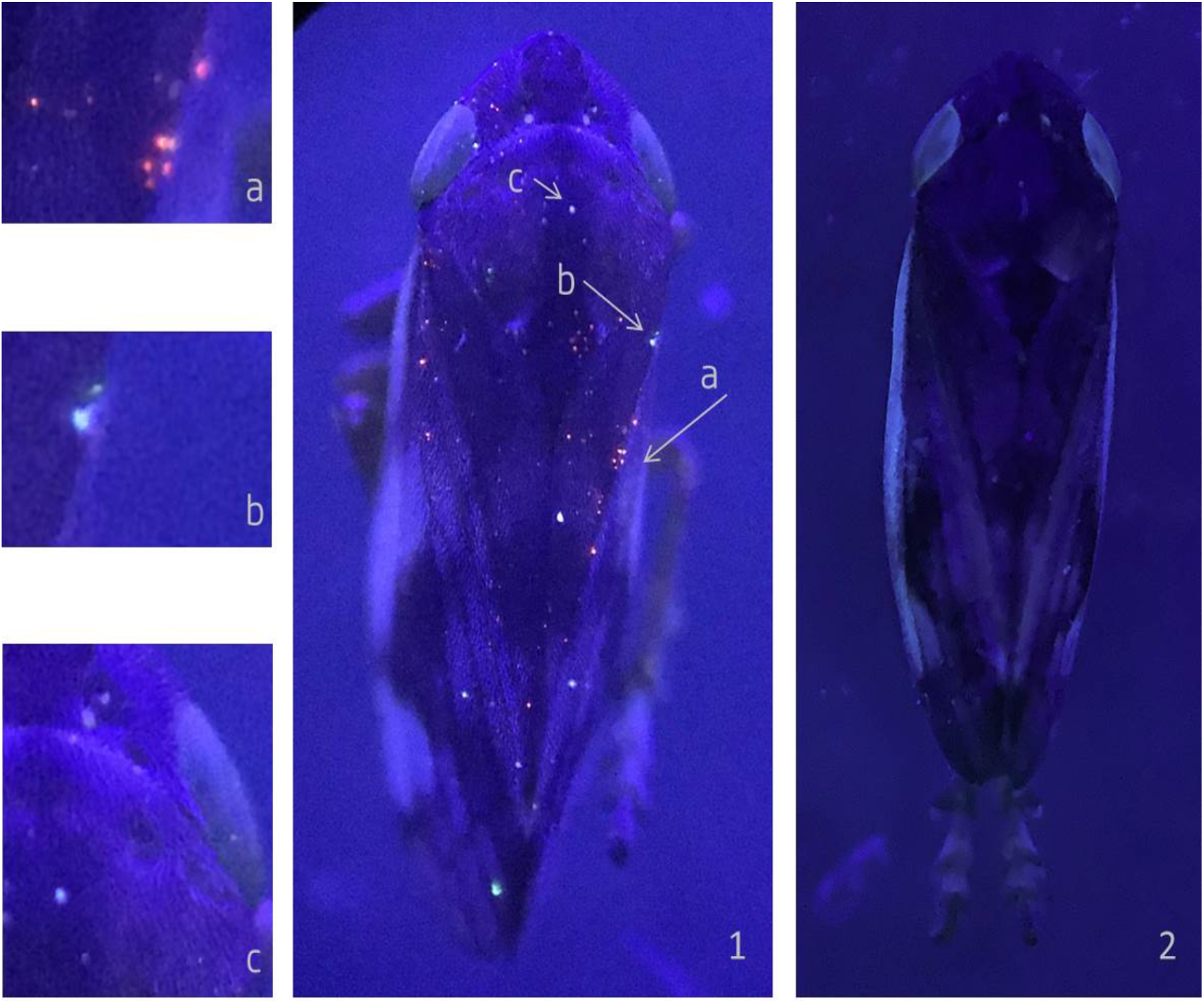
(1) An orange marked *N. campestris* recaptured in the zone D and exposed to UV light. Orange fluorescent particles were clearly visible (a). Other particles were found that could be either yellow fluorescent particles (b) or dust (c) covering some parts of the insect’s body. (2) An undusted individual of *N. campestris* exposed to UV light.

### 2.4. Directional movement of spittlebugs in an olive grove

The survey was carried out in 2019 from mid-May to late June, in a low-input olive grove located in “Villa del Prado” in the southwest of Madrid, Spain. The grove contained abundant ground cover vegetation until it naturally dried out in the summer. The percentage of fresh ground vegetation was 70% at the first sampling date in May 24^th^ and completely dried out in early June. A vineyard and three managed conventional tillage olive groves surrounded the field of study. On one of the borders between the field of study and the surrounding olive groves there were a few number of *Quercus ilex* subsp. *ballota* trees (Figure 3).

**FIGURE 3.**
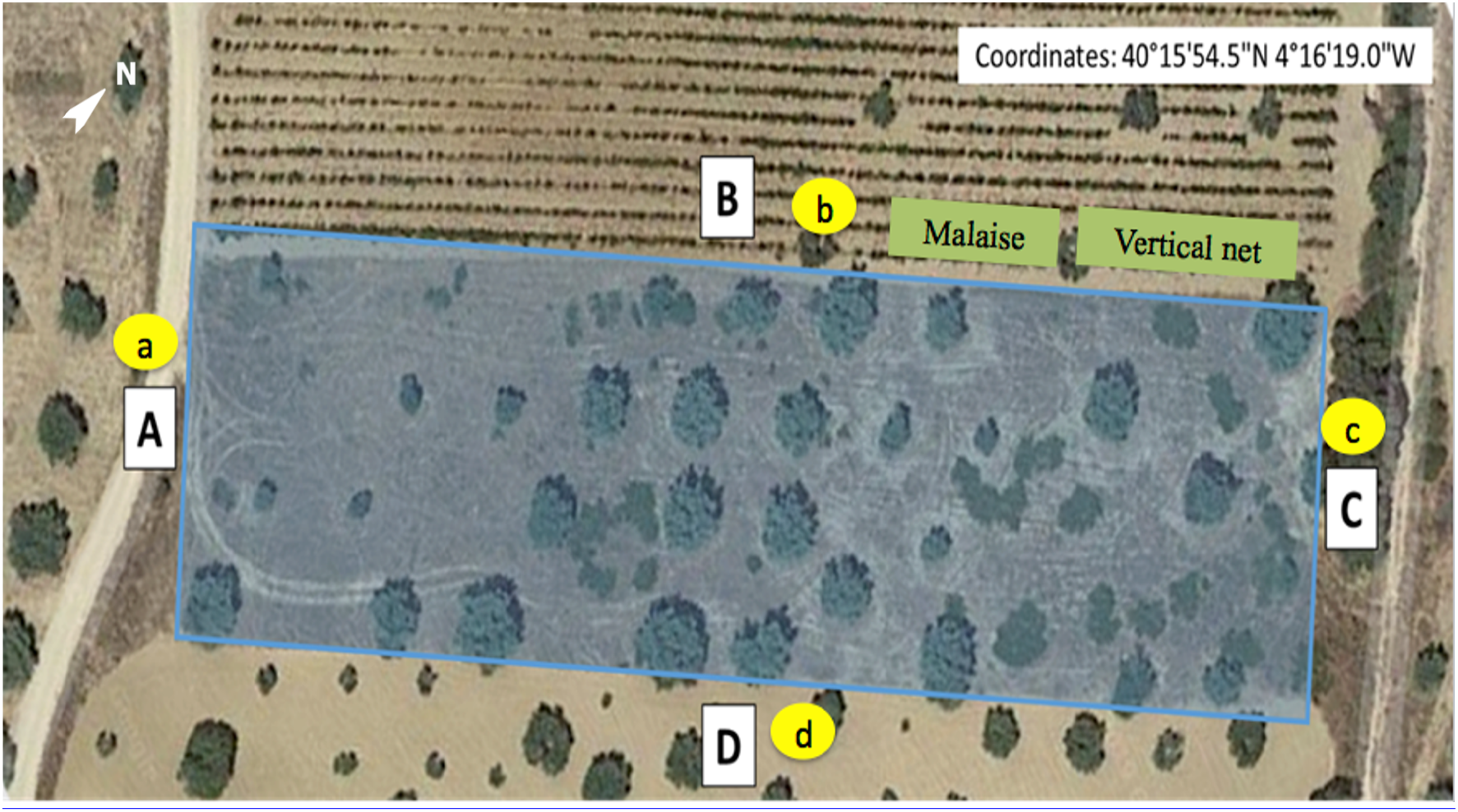
Olive grove in Villa del Prado. Size: 6.075 m^2^. The olive grove has 4 borders (A, B, C, D), surrounded by: (A) olive grove, (B) vineyard, (C) olive grove and a few *Q. ilex* subsp. *ballota* trees, (D) olive grove. One yellow sticky band was placed on each border (bands: a, b, c, d). A directional Malaise trap and a vertical net were also placed on the border (B). Google. (n.d.). Villa del prado. Retrieved from: https://www.google.es/maps/place/40%C2%B015'54.5%22N+4%C2%B016’19.0%22W/@40.2651569,-4.272151,132m/data=!3m1!1e3!4m5!3m4!1s0x0:0x0!8m2!3d40.2651389!4d-4.2719444?hl=es

Adults of putative vectors of *X. fastidiosa* (Hemiptera: Cercopoidea) were sampled by using 4 vertical yellow sticky traps, 1 directional Malaise trap and 1 vertical yellow sticky net.

The 4 vertical yellow sticky traps (Econex 100 m × 30 cm TA123, Murcia, Spain) were composed of two sticky bands with plastic surfaces (Figure 4). The bands were held with two steel sticks placed 170 cm apart from each other. The two bands were placed at two different heights: one at 20 cm and the other at 130 cm above the ground (Figure 4).

**FIGURE 4.**
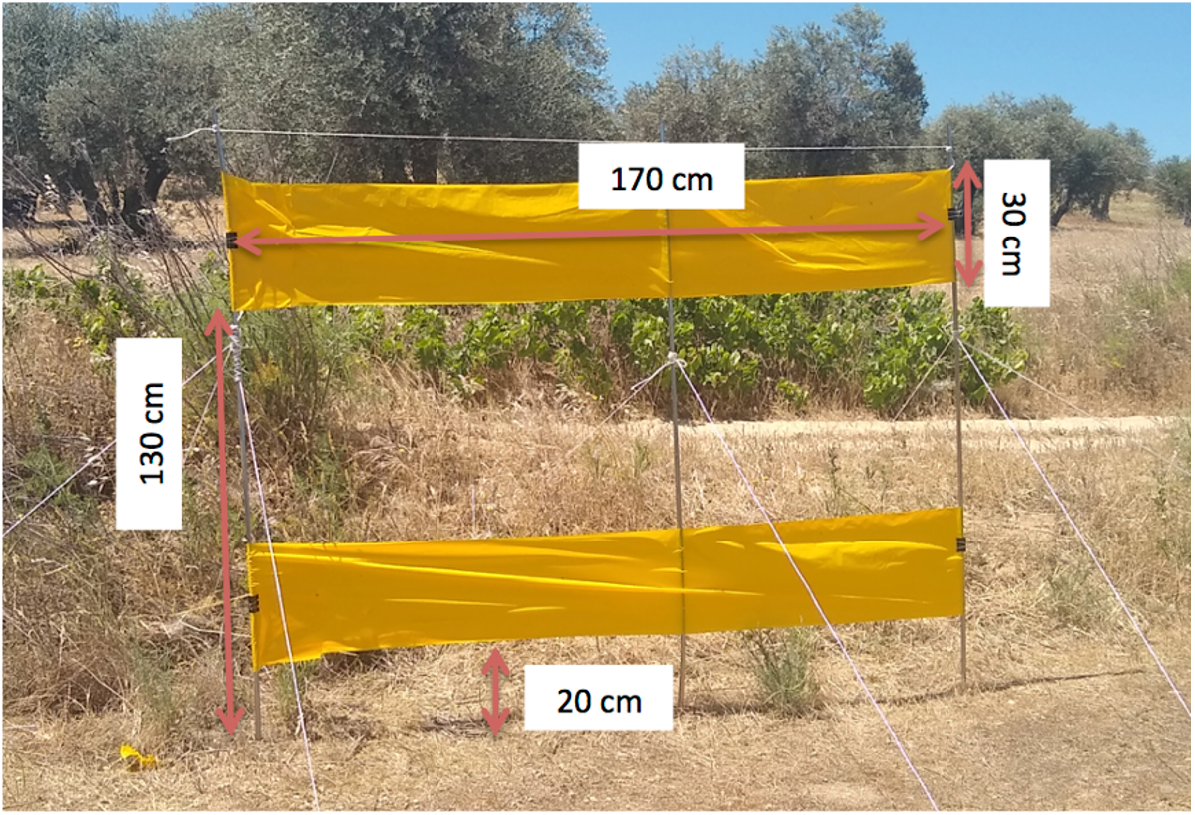
One of the 4 vertical yellow sticky trap used in the survey of directional movement on potential vectors of *X. fastidiosa* in the olive grove placed in Villa del Prado, Madrid, Spain.

The Malaise trap (BT1003, MegaView Science Co., Ltd., Taichung, Taiwan) used has two collecting heads with bottles filled with glycerol (50%) for directional sampling. Each bottle contained insects intercepted by each side of the vertical net. Therefore, catches represent insects emigrating or immigrating out or into the field of study.

The vertical yellow net (size: 2×3m; mesh: 7×7 threads/cm) was sprayed with glue (Souverode aerosol, Plantin SARL, Courthézon, France) to facilitate catches.

The four yellow sticky traps were placed between the field of study and the surrounding fields, one in each of the borders of the olive grove (Figure 3: A, B, C, D), facing each of the two sides of the olive grove, allowing directional sampling to differentiate between immigration and emigration out and into the field. The 4 yellow sticky traps were checked and replaced once a week. The Malaise trap and the vertical net trap were placed between the vineyard and the olive grove of study once a week (Figure 3: B). We checked the latter traps every half an hour from 9AM to 2PM and then they were removed. All the spittlebugs captured were counted and identified in the field. We kept track of insect directional movement by counting insects trapped on both sides of each type of trap (those emigrating and immigrating out or into the field of study).

## 3. RESULTS

### 3.1. Persistence of fluorescent dusts and their effects on survival and flight activity of *Neophilaenus campestris*

#### 3.1.1. Persistence of dusts and effects on N. campestris survival

Dusted and undusted *N. campestris* maintained under greenhouse conditions did not present significant differences in survival (two-sample Cox proportional hazards model Z= 1.271, P= 0.204) (Figure 5). Moreover, none of the marked individuals manifested a loss of marking dust beyond the level 2 during the 35 days’ period of the experiment being all the marked insects easily distinguishable under a naked eye. It is worth noting that the indoor environmental conditions where the insects were raised were different from those in the field. Insects were maintained inside cages in a glasshouse with no exposure to wind, rain or strong UV radiation.

**FIGURE 5.**
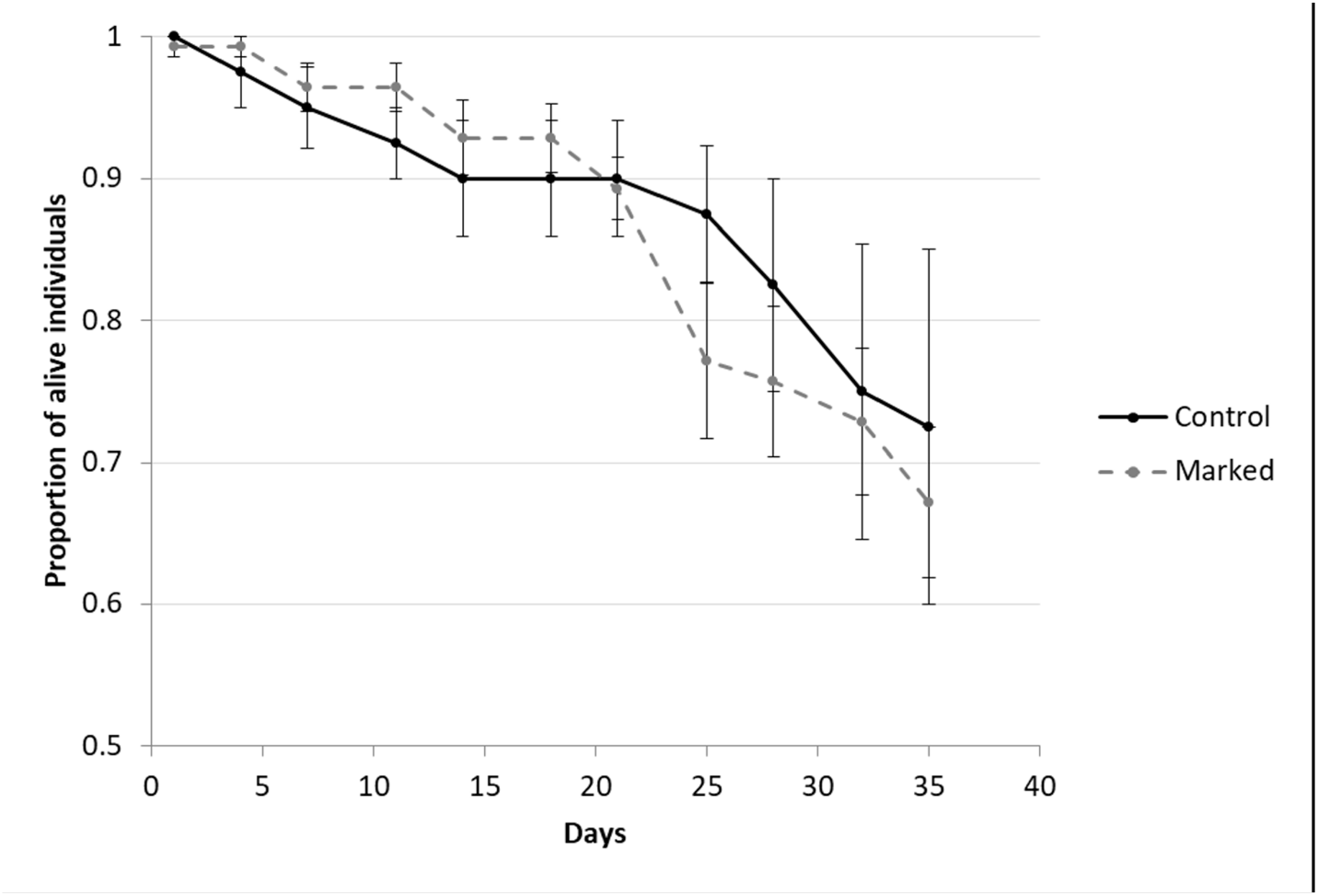
Survivorship curves for insects marked with fluorescence dust (dashed line) and undusted control insects (continuous line). Standard error bars are shown.

### 3.1.2. Dust effect on the flight activity of N. campestris

Flight mill assays showed that the overall proportion of individuals of *N. campestris* that were able to fly was 56.2% (50/89). There were no significant differences (df= 4; Chi-2 = 1.913; P= 0.752) between dusted and undusted insects in the proportion of individuals able to fly. Furthermore, no significant differences were found between dusted and undusted individuals for any of the flight descriptors measured: number of flights, total distance travelled, total duration of flight and average speed of flight (Table 1), according to ANOVA or Kruskal-Wallis tests. Therefore, all the data of dusted and undusted individuals was pulled together and the flight descriptors were calculated for all insects (n=50) (Table 2). *N. campestris* travelled 282 m in about 17 min in average in a single flight, and one individual was able to travel almost 1.4 km in an 82 minutes’ single flight. The mean speed of flight was 0.26 m/s.

**TABLE 1.**
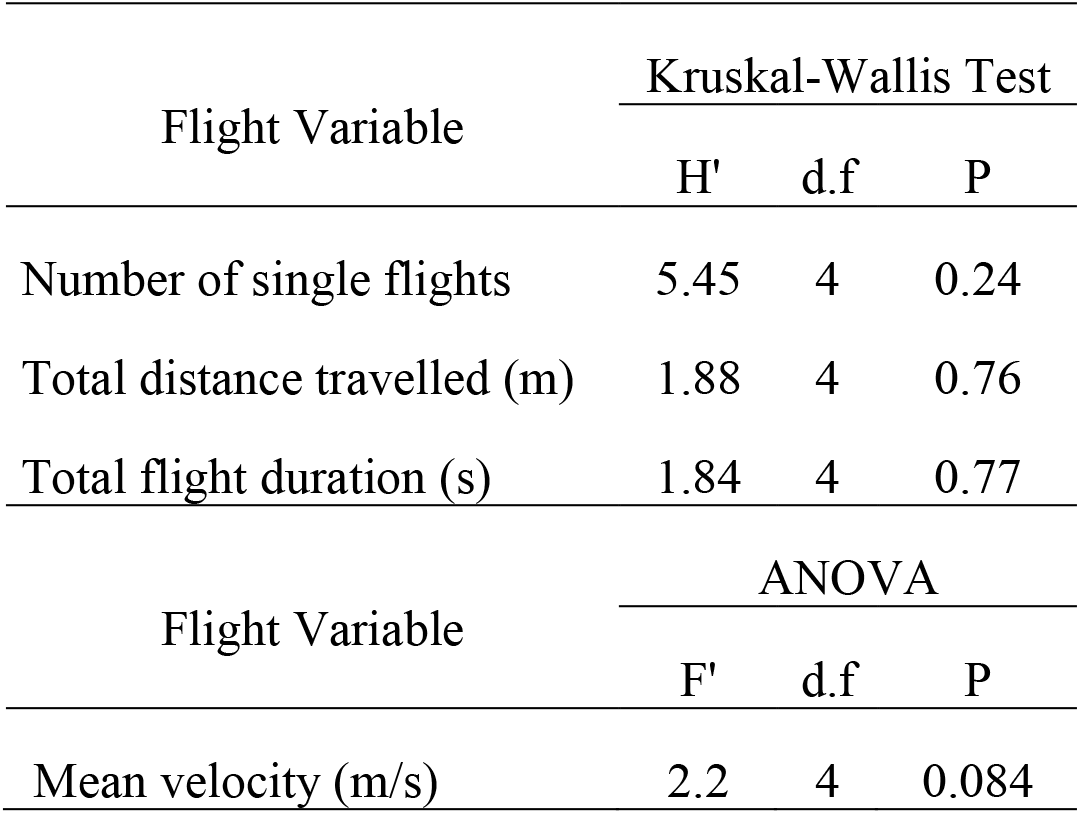
Summary of ANOVA or Kruskal-Wallis Test (selected depending on the data distribution) comparing dusted (pink, blue, yellow, orange; n=10 marked insects for each colour) and undusted (n=10) *N. campestris* adults (n=50).

| Flight Variable | Kruskal-Wallis Test |  |  |
| --- | --- | --- | --- |
|  | H' | d.f | P |
| Number of single flights | 5.45 | 4 | 0.24 |
| Total distance travelled (m) | 1.88 | 4 | 0.76 |
| Total flight duration (s) | 1.84 | 4 | 0.77 |
| Flight Variable | ANOVA |  |  |
|  | F' | d.f | P |
| Mean velocity (m/s) | 2.2 | 4 | 0.084 |

**TABLE 2.** Flight descriptors data. Results are showed as total values (mean ± SE) and maximum values in a single flight of *N. campestris* adults (n=50).

| Flight descriptors | Total (mean ± SE) | Maximum |
| --- | --- | --- |
| Number of single flights | 2.1 ± 0.26 | 10 |
| Distance travelled (m) | 281.54 ± 40.53 | 1362.3 |
| Flight duration (sec) | 1012.92 ± 137.62 | 4933.14 (≈82 min) |
| Flight speed (m/s) | 0.26 ± 0.01 | 0.42 |

### 3.2. Mass-Mark-Recapture assay (MMR)

During the MRR assay (23^rd^ May – 5^th^ July) the temperatures averaged 23.4 ± 0.78°C and the wind speed varied over the course of the day with a mean of 2.5 ± 0.14 m/s showing rates of maximum wind speed of 4.2 m/s and minimum of 1.8 m/s. A total of 1315 individuals of *N. campestris*, 430 individuals of *Lepyronia coleoptrata* and 30 individuals of *P.spumarius* were released. All the marked individuals that were recaptured were *N. campestris* and we were unable to recover any marked *P. spumarius* or *L. coleoptrata* in the sampling sites. Therefore, the results refer exclusively to *N. campestris* (Table 3). From a total of 1315 marked and released *N. campestris* (from the four dusted groups), 21 marked individuals were recaptured representing a mark-recapture rate of 1.6% (Table 3). A total of 791 individuals of *N. campestris* (considering both marked individuals and “wild” not marked insects) were captured from the 12 recapture sampling sites. However, recaptures of marked individuals occurred only in three (D, G and K) of the 12 sites sampled (Figure 1). The marked individuals that were recaptured were found only on two different species of pine trees: *P. halepensis* and *P. pinea*.

**TABLE 3.** Number of individuals of *Neophilaenus campestris* recaptured. The “Total Recaptured” column shows the total number of individuals of *N. campestris* (dusted and undusted) captured in the recapture zones. The ratios are expressed in parts per unit. Description and location of the release and recapture sites is shown in Figure. 1.

|  |  |  | No. marked per colour |  |  |  | No. marked / No. released per colour |  |  |  |
| --- | --- | --- | --- | --- | --- | --- | --- | --- | --- | --- |
| Zone | No. released | No. recaptured | Pink | Blue | Yellow | Orange | Pink | Blue | Yellow | Orange |
| Release pink | 335 | 0 | 0 | 0 | 0 | 0 | 0 | 0 | 0 | 0 |
| Release blue | 350 | 0 | 0 | 0 | 0 | 0 | 0 | 0 | 0 | 0 |
| Release yellow | 300 | 0 | 0 | 0 | 0 | 0 | 0 | 0 | 0 | 0 |
| Release orange | 330 | 0 | 0 | 0 | 0 | 0 | 0 | 0 | 0 | 0 |
| A | - | 0 | 0 | 0 | 0 | 0 | 0 | 0 | 0 | 0 |
| B | - | 0 | 0 | 0 | 0 | 0 | 0 | 0 | 0 | 0 |
| C | - | 0 | 0 | 0 | 0 | 0 | 0 | 0 | 0 | 0 |
| D | - | 209 | 0 | 0 | 0 | 8 | 0 | 0 | 0 | ≈0.02 |
| E | - | 0 | 0 | 0 | 0 | 0 | 0 | 0 | 0 | 0 |
| F | - | 0 | 0 | 0 | 0 | 0 | 0 | 0 | 0 | 0 |
| G | - | 107 | 0 | 0 | 0 | 8 | 0 | 0 | 0 | ≈0.02 |
| H | - | 8 | 0 | 0 | 0 | 0 | 0 | 0 | 0 | 0 |
| I | - | 73 | 0 | 0 | 0 | 0 | 0 | 0 | 0 | 0 |
| J | - | 0 | 0 | 0 | 0 | 0 | 0 | 0 | 0 | 0 |
| K | - | 402 | 0 | 0 | 1 | 4 | 0 | 0 | 0.003 | ≈0.01 |
| L | - | 0 | 0 | 0 | 0 | 0 | 0 | 0 | 0 | 0 |
| Total | 1315 | 791 | 0 | 0 | 1 | 20 | 0 | 0 | 0.003 | ≈0.05 |

All the individuals recaptured were dusted with either orange or yellow dusts. No individuals with a blue or pink dust were recaptured.

*N. campestris* recaptured in points D (8 individuals) and G (8 individuals) were marked with the orange colour (Figure 2) which indicated that these insects flew 123 m from the orange release point to the D zone and 281 m to the G zone. Furthermore, 5 dusted individuals of *N. campestris* were recaptured in the K point, which was about 2000 meters away from the release point. Four of these 5 individuals presented orange dust while 1 individual was marked only with yellow fluorescent dust.

The majority of the orange dusted insects presented many orange dots and few yellow or whitish dust particles on their body (Figure. 2) but for the purpose of the analysis we considered all marked individuals in zones D and G coming from the orange release site (Table 2).

In point D, recaptures were possible throughout the whole assay. Thus, 3 orange-marked *N. campestris* were recaptured on 12^th^ June, 3 on 19^th^ June and 2 individuals on 27^th^ June. By contrast, in point G the only date of recapture was 5^th^ July when the 8 orange-marked *N. campestris* were recaptured. Finally, in the point K, the 4 orange-marked and the full yellow-marked *N. campestris* were captured on 27^th^ June. Recaptures in point D were done under variable climatic conditions while in point G and point K, recaptures matched with the two of the most windy and hottest days of the recapturing period : 27.94 °C of temperature and 2.9 m/s of wind speed and 30.96°C and 4.15 m/s respectively.

### 3.3. Directional movement of spittlebugs in an olive grove

Most putative vectors of *X. fastidiosa* collected in our survey were caught in the yellow sticky bands. Thus, we collected the greatest number of vectors in band b (Figure 3), situated between the vineyard and the olive grove of study. Results indicated a greater number of immigrating insects coming from the vineyard to the olive grove: 26 in total, 10 *N. campestris*; nine *P. spumarius*; three. *L. coleoptrata*; four *Cercopis spp*. The number of potential vectors emigrating from the olive grove to the vineyard was lower: 16 in total, eight *N. campestris*; three *P. spumarius*; one *L. coleoptrata*; four *Cercopis spp*. Most individuals were captured before second week of June (Figure 6), when the ground vegetation dried out. The band a (Figure 3), collected three immigrating spittlebugs (one *N. campestris* and two *P. spumarius*), while five emigrating spittlebugs (two *N. campestris*; three *P. spumarius*). The band c (Figure 3) collected one immigrating spittlebug (one *N. campestris*) and two emigrating spittlebugs, (one *N. campestris* one *P. spumarius*). Finally, in band d (Figure 3) one single *L. coleoptrata* and one *N. campestris* were captured immigrating and emigrating from the field of study, respectively. Regarding the Malaise and vertical net traps, there were almost no captures. A single *Lepyronia sp*. was captured with the vertical net and no potential vectors of *X. fastidiosa* were caught in the Malaise trap.

**FIGURE 6.**
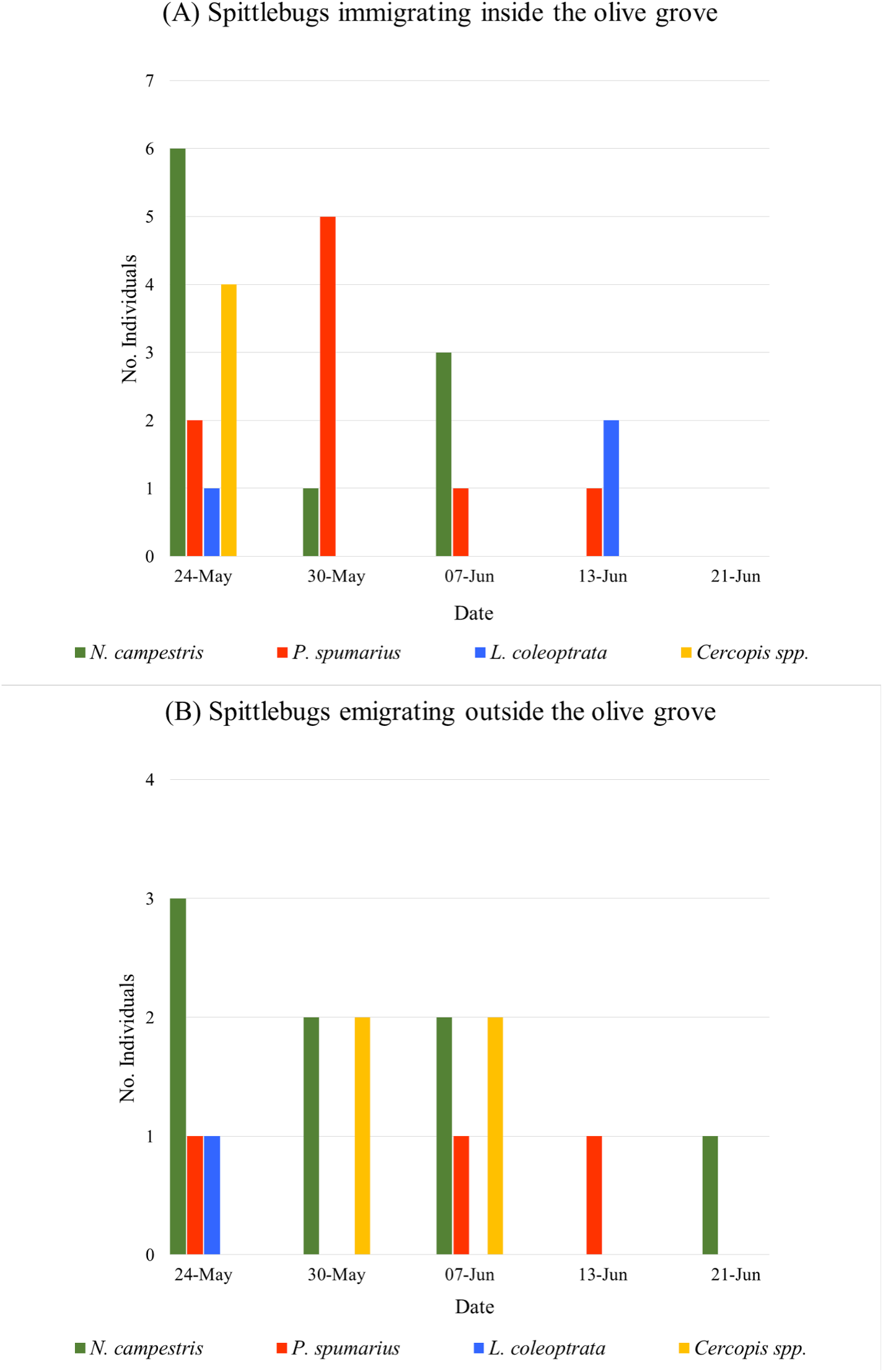
Number of spittlebug adults caught on the yellow sticky trap between the vineyard and the olive grove (Figure 3. trap b). (A) Spittlebugs immigrating from the vineyard to the olive grove. (B) Spittlebugs emigrating from the olive grove to the vineyard.

## 4. DISCUSSION

*N. campestris* is a vector of *X. fastidiosa* (Cavalieri et al., 2019) and has a widespread distribution across the Iberian Peninsula (Morente et al., 2018a). *N. campestris* spends most of its life cycle on the ground cover vegetation, mainly on grasses where mating, oviposition and feeding occur. However, this species moves from the ground cover to trees and shrubs in the late spring (Lopes et al 2014; Cornara et al., 2016; Morente et al., 2018a; Antonatos et al., 2019). To date, politics of containment, common to the European Union, relies on scarce information about the Cercopoidea dispersal abilities, most of them collected in landscapes different from the Mediterranean scrubland (Weaver & King, 1954; Halkka et al., 1971; Plazio et al., 2017). However, landscape composition and climate conditions can influence the distribution and movement of insects affecting the speed and the track of movement (Crist et al., 1992; Jonsen & Taylor, 2000, Haynes & Cronin, 2003, 2006; Blackmer et al., 2006). Thus, information about the movement ability of *N. campestris* or any other vector species in the landscape of interest (an olive grove and the natural Mediterranean landscape surrounding in the present study) can be crucial to adopt effective solutions to contain the spread of *X. fastidiosa* in Europe.

Our indoor tests on survival, dust retention and flying capabilities of *N. campestris* showed that the methodology applied in our MRR field assay did not disturb the flight behavior or survival of the dusted spittlebugs. However, insects exposed to natural conditions were different to those exposed to indoor facilities as they are protected from rain and intensive UV light. This could explain why the marked insects collected in the field were not visible under naked eye and a UV lamp was always needed for detection of the fluorescent dust.

The flight mill study also showed that the flight potential of *N. campestris* was much higher than was previously assumed. Flight mill data can be difficult to interpret because insect’s behavior and flying ability could be influenced because of manipulation. However, flight mill experiments allow comparing differences in flight behaviour between different groups, such as different insect species, ages, sexes or mated status (Dingle, 1966; Avalos et al., 2014; Minter et al., 2018; Guo et al., 2020). In the flight mill assay we found that dust marking did not affect the flight potential of *N. campestris*. Moreover, this assay estimates the flight potential of these insects, showing that they are able to travel much more than 100 m in less than an hour.

The results obtained in the MMR assay support previous studies (Lopes et al., 2014 and Morente et al., 2018a), which proved that *N. campestris* move and settle on pine trees during late spring and summer (in our study *P. pinea* and *P. halepensis*). Therefore, the spittlebugs that were recaptured in the K zone were able to travel distances longer than 2 km. Those that came from the orange release point travelled around 2282 m and those that came from the yellow release point moved a total of 2473 m, the longest distance covered by a spittlebug recorded in a field assay until now (Freeman, 1945; Weaver & King, 1954; Reynolds et al., 2017). Thus, our results suggest that *N. campestris* is able to travel more than 2000 meters in 35 days. As other spittlebugs, the long-distance movement of *N. campestris* could be dependent on the winds. Thus, *P. spumarius* or *N. lineatus* might be capable to fly some meters up reaching the air currents to migrate passively (Freeman, 1945; Weaver & King, 1954; Reynolds et al., 2017). Regarding short-distance migration, we can assert that *N. campestris* is able to move more than 100 m in 24 hours. The changing wind conditions during the assay did not enable us to identify a dominant wind pattern. Therefore, other variables such as the presence of resting places in the migration track, that serve as corridor to sheltered places, may favor the migration of *N. campestris* (Hunter, 2002). Moreover, most orange dusted insects presented some yellow spots on their body. However, except in the K point, where we found a yellow spittlebug, we did not find any other yellow dusted *N. campestris* in other zones. Mixed patches of olive groves and pine and oak woods surrounded the yellow and orange release points. Yellow and orange dusted *N. campestris* could have met in a middle resting point within the migration track where they could mate and thus transferred the dust from one individual to another. More likely, yellow dusted insects may have transferred some dust particles to the orange-marked insects while they remained in the cage in the laboratory before sorting them out in the microscope. When the recaptured insects were placed in the falcon tubes and cages they were able to contact each other and mate. Therefore, transfer of dust from one insect to another is a possibility that cannot be excluded. Finally, no *N. campestris* was found on the rest of oversummering host plants sampled in the study such as oak trees. This result may indicate that, despite the polyphagous character of the insect, it presents a strong migratory preference for pines in the summer. Morente et al., 2018a described that *N. campestris* tend to return back to olive groves in the fall to lay the eggs. Thus, the presence of pines in the landscape surrounding the crop may favor the establishment and proliferation of *N. campestris* in a given area throughout the year. Thus, nymphs grow up on the ground cover-mainly grasses-in olive groves and the emerged adults spend most of the summer on the surrounding pine trees returning to the olive grove after the first rains in the fall to mate and lay their eggs on the emerging grasses. This has been observed in several areas of Spain including the Alicante region where *N. campestris* was very abundant in the grasses during the fall (Morente et al., 2018b)

In the survey of directional movement on potential vectors of *X. fastidiosa*, most of the spittlebugs were captured in the yellow sticky band placed between the olive grove and the vineyard. In contrast, there were fewer captures in the other three yellow sticky bands, placed between the olive grove of study and the other three olive groves. Moreover, few spittlebugs were captured after the ground vegetation dried out (Figure 6). This can be explained because of the lack of weeds in the surrounding olive groves that contrast with the succulence of the grapevine leaves and the presence of herbaceous vegetation cover in our field of study. A seasonal pattern in spittlebugs movement depending on the ground vegetation cover has been previously observed by several authors (Weaver & King, 1954; Waloff, 1973; Nilakhe & Buainain, 1988; Cornara et al., 2017a; Cruaud et al., 2018; Cornara et al., 2018; Bodino et al. 2019). This suggests that these insects movement depends on the succulence of the plants available, so the non-tillage practices, and the presence of succulent plants in an area, could enhance the *X. fastidiosa* spread between agroecosystems. Also, the lack of catches in the Malaise trap and the vertical net in contrast to the yellow sticky bands, suggest that they are not a suitable sampling method for studying directional movement of spittlebugs. Other methods that allow a higher number of catches should be developed to study movement and migration of spittlebugs.

The short and long-distance migration capacity of *N. campestris* is added to a long list of difficulties that hamper the implementation of effective measurements of disease containment. Thus, our results showing that *N. campestris* can migrate and fly more than 2 km in 5 weeks together with the polyphagous habit of this species and its long life cycle invite to reconsider the methods applied until now in the infected crops. In fact, a recent modelling study by Strona et al (2020) shows that even limited probabilities of long distance dispersal of infectious vectors dramatically affected disease outbreaks caused by *X. fastidiosa* in olive groves in Andalusia (southern Spain). They concluded that identifying (and disrupting) long distance dispersal processes may be much more effective to contain disease epidemics than surveillance and intervention concentrated on local scale transmission processes. Thus, the eradication measures that are being adopted as a general rule in the EU to fight against the disease by up-rooting the infected plants and the entire host plants in a radius of 100 m, regardless their health status might not be the best strategy to contain the spread of diseases caused by *X. fastidiosa*. The fact that vectors of *X. fastidiosa* are able to fly much more than 100 m, the persistence of the disease in the vector for their entire life (almost 1 year) together with the polyphagous nature of most xylem-feeders suggests that up-rooting uninfected trees in infected areas-may have a limited impact in containing the disease.

*X. fastidiosa* symptoms onset is variable depending on the plant species, from three to four months (as in grapevine) to years (in the case of the olive tree) (Almeida 2016). Moreover, the detection of the pathogen within the plant is a difficult task, which requires certain concentrations of the bacterium and the right collection of samples (at least 4 leaves per sample from different trees in a large scale sampling in the case of olive tree) (Loconsole et al., 2014). Additionally, vectors may acquire *X. fastidiosa* from infected asymptomatic hosts. Another important finding is that transmission of *X. fastidiosa* by their vectors is a very fast process (inoculation occurs in 2 to 7 minutes after the onset of the first probe) (Cornara et al., 2020a). Therefore, despite the fact that *N. campestris* adults do not colonize olive trees (Mazzoni, 2005; Cornara et al. 2017a; Morente et al., 2018a; Bodino et al., 2019) they could easily land and probe briefly on olive canopies in late spring and summer when they disperse towards their over-summering hosts. In this process they could rapidly transmit *X. fastidiosa* from one tree to another. Thus, as suggested by Almeida (2016) eliminating the trees in a delimited infected area may have a limited impact in the disease containment. In summary, the main strategy to contain and limit *X. fastidiosa* outbreaks should be focused in managing vector populations in their early stage of development (immature stages). This will avoid or reduce the presence of adults in areas were the disease is present and limit the risk of long distance dispersal. This goal should be addressed in the most sustainable way by understanding the ecology, biology and behaviour of spittlebugs. Cultural control tactics such as conservation tillage in the right moment will be essential to disrupt the life cycle of spittlebugs.

Finally, our study concentrated on *N. campestris,* which was the most abundant vector species in our area of study. However, the main vector of *X. fastidiosa* in Europe is *P. spumarius* (Cornara et al., 2019), thus additional studies are needed to determine the migration behaviour of *P. spumarius* in areas where this vector is the dominant species.

## Supporting information

Supplemental material Annex 1

Supplemental material Graphical_Abstract

## 5. ACKNOWLEDGMENTS

Authors would like to acknowledge our colleagues Marcos Ramirez, Joao Leitao and Pierpaolo Natilla for their help in the field work. The work has been funded by the Ministerio de Ciencia e Innovación under the grant agreement no. AGL2017-89604-R and by Comunidad de Madrid under the grant agreement FP19-XYLELLA: Evaluación de la incidencia de *Xylella fastidiosa* en la Comunidad de Madrid.

## 6 TABLES & FIGURES

**ANNEX 1.** Distance (m.) between the spittlebug release (colours) and the recapture (letters) points.

