## Supplemental material Annex 1 for "Dispersal ability of *Neophilaenus campestris*, a vector of *Xylella fastidiosa*, from olive groves to over-summering hosts"

**ANNEX 1.** Distance (m.) between the release (colours) and the recapture (letters) points.

| | Pink | Blue | Yellow | Orange |
|---|---|---|---|---|
| **Pink** | 0.00 | 183.70 | 316.91 | 366.30 |
| **Blue** | 183.70 | 0.00 | 193.92 | 375.35 |
| **Yellow** | 316.91 | 193.92 | 0.00 | 205.92 |
| **Orange** | 366.30 | 375.35 | 205.92 | 0.00 |
| **A** | 412.27 | 506.95 | 430.09 | 614.29 |
| **B** | 94.26 | 209.27 | 248.58 | 477.76 |
| **C** | 575.89 | 477.42 | 277.12 | 139.01 |
| **D** | 640.90 | 490.55 | 323.15 | 123.00 |
| **E** | 788.85 | 685.86 | 491.76 | 338.86 |
| **F** | 762.45 | 637.46 | 462.13 | 261.22 |
| **G** | 791.26 | 637.25 | 494.84 | 280.85 |
| **H** | 1034.07 | 850.59 | 729.75 | 515.27 |
| **I** | 1409.81 | 1243.39 | 1109.13 | 870.07 |
| **J** | 1534.70 | 1356.83 | 1303.54 | 1133.27 |
| **K** | 2753.61 | 2564.81 | 2472.77 | 2282.51 |
| **L** | 1741.18 | 1900.61 | 2021.92 | 2252.13 |
