## Supplementary figures and images for "Dispersal ability of *Neophilaenus campestris*, a vector of *Xylella fastidiosa*, from olive groves to over-summering hosts"

### Supplemental material Graphical_Abstract

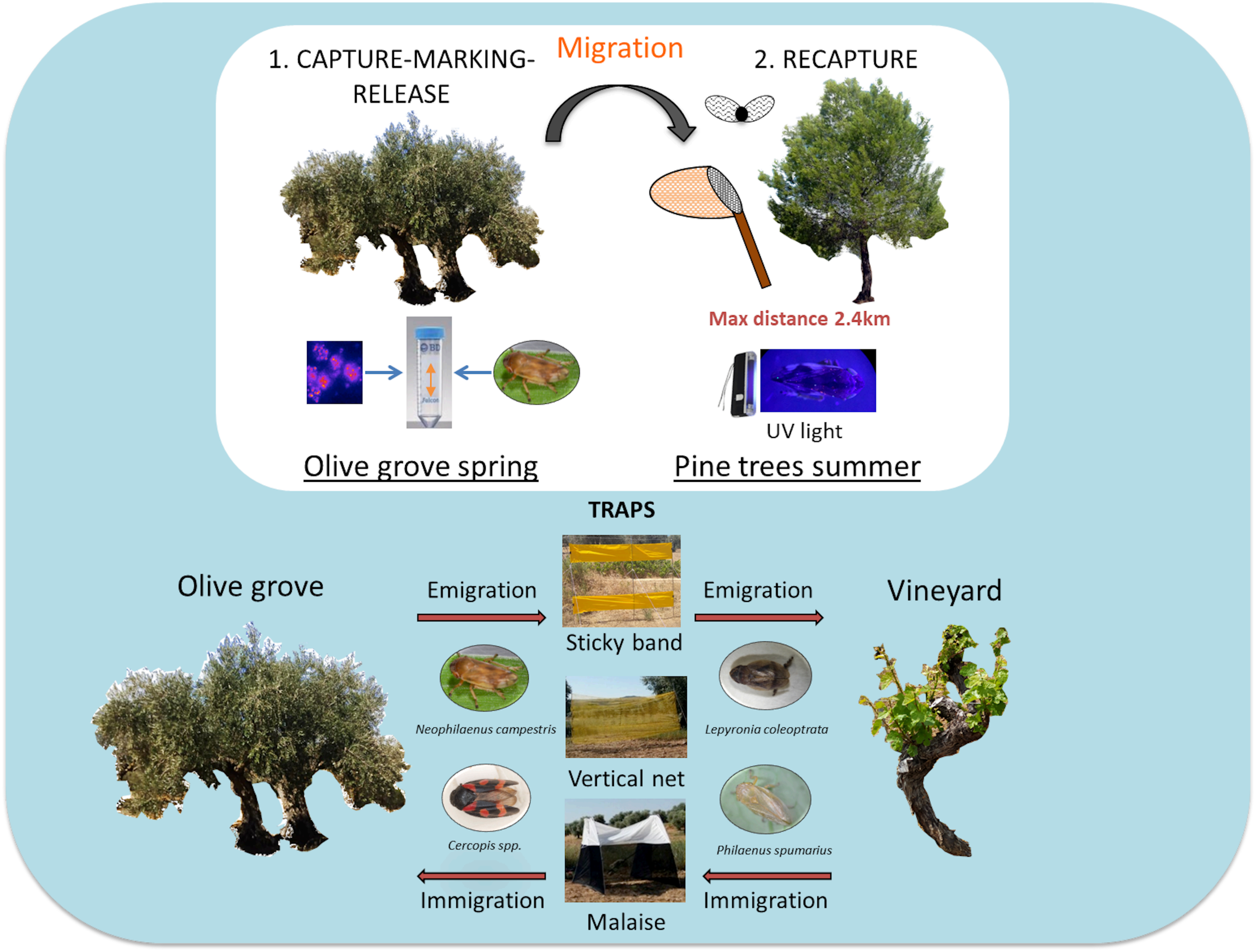
